# Analysis of putative cis-regulatory elements regulating blood pressure variation

**DOI:** 10.1101/820522

**Authors:** Priyanka Nandakumar, Dongwon Lee, Thomas J. Hoffmann, Georg B. Ehret, Dan Arking, Dilrini Ranatunga, Man Li, Megan L. Grove, Eric Boerwinkle, Catherine Schaefer, Pui-Yan Kwok, Carlos Iribarren, Neil Risch, Aravinda Chakravarti

## Abstract

Hundreds of loci have been associated with blood pressure traits from many genome-wide association studies. We identified an enrichment of these loci in aorta and tibial artery expression quantitative trait loci in our previous work in ∼100,000 Genetic Epidemiology Research on Aging (GERA) study participants. In the present study, we subsequently focused on determining putative regulatory regions for these and other tissues of relevance to blood pressure, to both fine-map these loci by pinpointing genes and variants of functional interest within them, and to identify any novel genes.

We constructed maps of putative cis-regulatory elements using publicly available open chromatin data for the heart, aorta and tibial arteries, and multiple kidney cell types. Sequence variants within these regions may be evaluated quantitatively for their tissue- or cell-type-specific regulatory impact using deltaSVM functional scores, as described in our previous work. In order to identify genes of interest, we aggregate these variants in these putative cis-regulatory elements within 50Kb of the start or end of genes considered as “expressed” in these tissues or cell types using publicly available gene expression data, and use the deltaSVM scores as weights in the well-known group-wise sequence kernel association test (SKAT). We test for association with both blood pressure traits as well as expression within these tissues or cell types of interest, and identify several genes, including *MTHFR*, *C10orf32*, *CSK*, *NOV*, *ULK4*, *SDCCAG8*, *SCAMP5*, *RPP25*, *HDGFRP3*, *VPS37B*, and *PPCDC*. Although our study centers on blood pressure traits, we additionally examined two known genes, *SCN5A* and *NOS1AP* involved in the cardiac trait QT interval, in the Atherosclerosis Risk in Communities Study (ARIC), as a positive control, and observed an expected heart-specific effect. Thus, our method may be used to identify variants and genes for further functional testing using tissue- or cell-type-specific putative regulatory information.

**Author Summary:** Sequence change in genes (“variants”) are linked to the presence and severity of different traits or diseases. However, as genes may be expressed in different tissues and at different times and degrees, using this information is expected to more accurately identify genes of interest. Variants within the genes are essential, but also in the sequences (“regulatory elements”) that control the genes’ expression in different tissues or cell types. In this study, we aim to use this information about expression and variants potentially involved in gene expression regulation to better pinpoint genes and variants in regulatory elements of interest for blood pressure regulation. We do so by taking advantage of such data that are publicly available, and use methods to combine information about variants in aggregate within a gene’s putative regulatory elements in tissues thought to be relevant for blood pressure, and identify several genes, meant to enable experimental follow-up.

## Introduction

Genetic studies of complex disorders have identified hundreds to thousands of sequence variants in the human non-coding genome. However, despite significant mapping progress, we do not yet know the identity of most of the underlying genes and variants, nor have a mechanistic understanding of how these genes, individually and together, contribute to a phenotype. Thus, we need to consider how such genomic studies can improve our knowledge of trait physiology. One approach would be to focus study genetic analyses by organs and tissues of interest.

Pritchard and colleagues have hypothesized that the majority of genome-wide association study (GWAS) signals may be functionally spurious and arise from genes peripheral to the core functions affected in a trait or disease [1]. These false positives dominate because most genes in a cell-type are connected by gene expression to one another through very shallow functional networks, a working hypothesis that fails to explain the stability of network perturbations (robustness) or their specificity (phenotypic effects) [2–4]. To resolve this question, connecting genotypes to phenotypes through gene expression variation is of primary importance since eQTLs (expression quantitative trait loci) are identifiable causal factors [5, 6]. However, utilizing gene expression in trait-related tissues is necessary [7], as genes exert their activities in the context of a core genetic network with intrinsic (cell autonomous) and extrinsic (non-autonomous) feedback [8].

Transcription within mammalian genomes is locally regulated within ∼400 kilobase (kb) chromatin segments called topological associating domains (TADs), largely invariant across cell types [9]. TADs contain numerous dispersed spatiotemporal expression cis-regulatory elements (CREs or enhancers) that are are brought together by DNA looping to allow binding of various transcription factors (TF) to enable gene expression control [10]. Many enhancers are recognized by their DNaseI hyper-sensitivity (DHS), ATAC-seq (Assay for Transposase-Accessible Chromatin using sequencing) assays [11], or adjacent histone (H3K4me1, H3K4me3, H3K27ac) modifications [12, 13]. Their phenotypic importance is evident from the fact that only 2.6% of the genome comprises DHS and histone marks [14] but explains ∼30% of the heritability of traits [15]. Thus, trait variation is from sequence changes within TFs, their binding sites (TFBS) and CREs, all detectable through epigenomic marks in cell lines and tissues. In this study, we propose an approach wherein these types of epigenomic data are used to identify genes within a GWAS locus in tissues of interest.

The analyses we propose are enabled by numerous public genomic resources. The Encyclopedia of DNA Elements (ENCODE) Project (https://www.encodeproject.org/) has generated open chromatin, RNA and DNA sequencing, genotyping, and histone modification data, among other data types. The Genotype-Tissue Expression (GTEx) Project (https://www.gtexportal.org/) includes genotype and expression data across 53 tissues and is useful as a reference transcriptome and eQTL dataset. These public resources also enable the development of an annotation score, deltaSVM [16], in which the quantitative impact of a non-coding variant on tissue or cell type specific gene regulation is predicted, based on a reference training set of regulatory regions. In this study, we exemplify this reverse genetic approach by focusing on blood pressure (BP) and QT interval variation.

Although the roles of the kidney and adrenal gland are well established in blood pressure regulation and syndromes [17–19], our previous work in the Kaiser Permanente Research Program on Genes, Environment and Health (RPGEH) Genetic Epidemiology Research on Adult Health and Aging (GERA) [20, 21] study demonstrated that associated variants at BP GWAS loci were enriched in eQTLs specific to the aorta and tibial arteries. Expanding on this work in this study, we aimed to connect groups of proximal putative regulatory variants within and around each gene to both the gene’s expression and also to BP traits, inferring that the gene’s expression in a potentially relevant tissue affected the regulation of BP. To accomplish this, within each artery dataset, we identified putative CREs, and by extension, putative CRE variants, for every gene, and tested these variants in aggregate for association with BP in the GERA study, as well as with expression in the GTEx study. We used the sequence kernel association test (SKAT) [22] for these association analyses, with each variant weighted by their deltaSVM score, to up-weight variants with greater predicted effects on gene regulatory activity. We supplemented our expression analyses with the software MetaXcan [23] to test whether the predicted expression of genes in each individual could be associated with BP. Prior to the novel BP gene discovery analyses of tissue involvement, and as a positive control, we first examined genes for the cardiac trait QT interval for which there is strong functional evidence of primarily heart involvement, using data from the Atherosclerosis Risk in Communities (ARIC) [24, 25] study. Finally, we examined the effects of putative regulatory variation for monogenic BP syndrome genes, all known to be renal or adrenal disorders, in four available kidney cell types to test for a group effect on BP.

Our results demonstrate the feasibility of identifying BP genes by tissue, which we expect will facilitate more comprehensive functional analyses of BP genes and BP control mechanisms.

## Results

We conducted several tissue-specific analyses to identify tissues and genes of interest for BP regulation using the GERA study. We initially focused on identifying tissues relevant to BP GWAS loci, and subsequently expanded on this by using tissue-specific information to analyze putative CRE variation of genes in these tissues. The aim was to identify specific genes and variants of interest at these GWAS loci. We also studied putative regulatory variation at 20 monogenic syndromic hypertension and hypotension genes in several kidney cell types. To begin, our study also includes an analysis of QT interval as a positive control to demonstrate the identification of well-characterized genes for that trait.

### Partitioned heritability of BP

We examined heritability for SBP and DBP using 80,792 GERA EUR subjects with stratified LD score regression (LDSC) across several functional categories [15], to identify functional categories in which BP heritability was enriched. We found that the top-ranked enriched categories were enhancer-associated histone marks H3K27ac, H3K4me1, and the Hnisz “super-enhancer” category (Tables S1-S2). This is in accordance with a previous study in which BP heritability was determined to be mostly from within DNaseI hypersensitivity sites [26], and, taken together with the results of the eQTL enrichment analyses, supports the study of regulatory elements in specific tissues of interest for BP.

### Constructing CRE maps

With knowledge of tissues highly relevant to characterizing BP GWAS loci, our next aim was to test each gene’s putative cis-regulatory variation for association with both gene expression and BP, in a tissue-specific context. This is expected to assist in identifying novel genes of interest, as well as provide tissue- or cell-type-specific information about known genes.

We first constructed CRE maps for the aorta and tibial arteries, as well as four kidney cell types (renal cortical epithelial cell, glomerular endothelial cell, epithelial cell of proximal tubule, and glomerular visceral epithelial cell), because of the known involvement of the kidney in blood pressure regulation [17, 18], using ENCODE data (Table 1) (though many monogenic forms of blood pressure disorders occur due to an effect of the adrenal gland on renal function [19]).

**Table 1.**
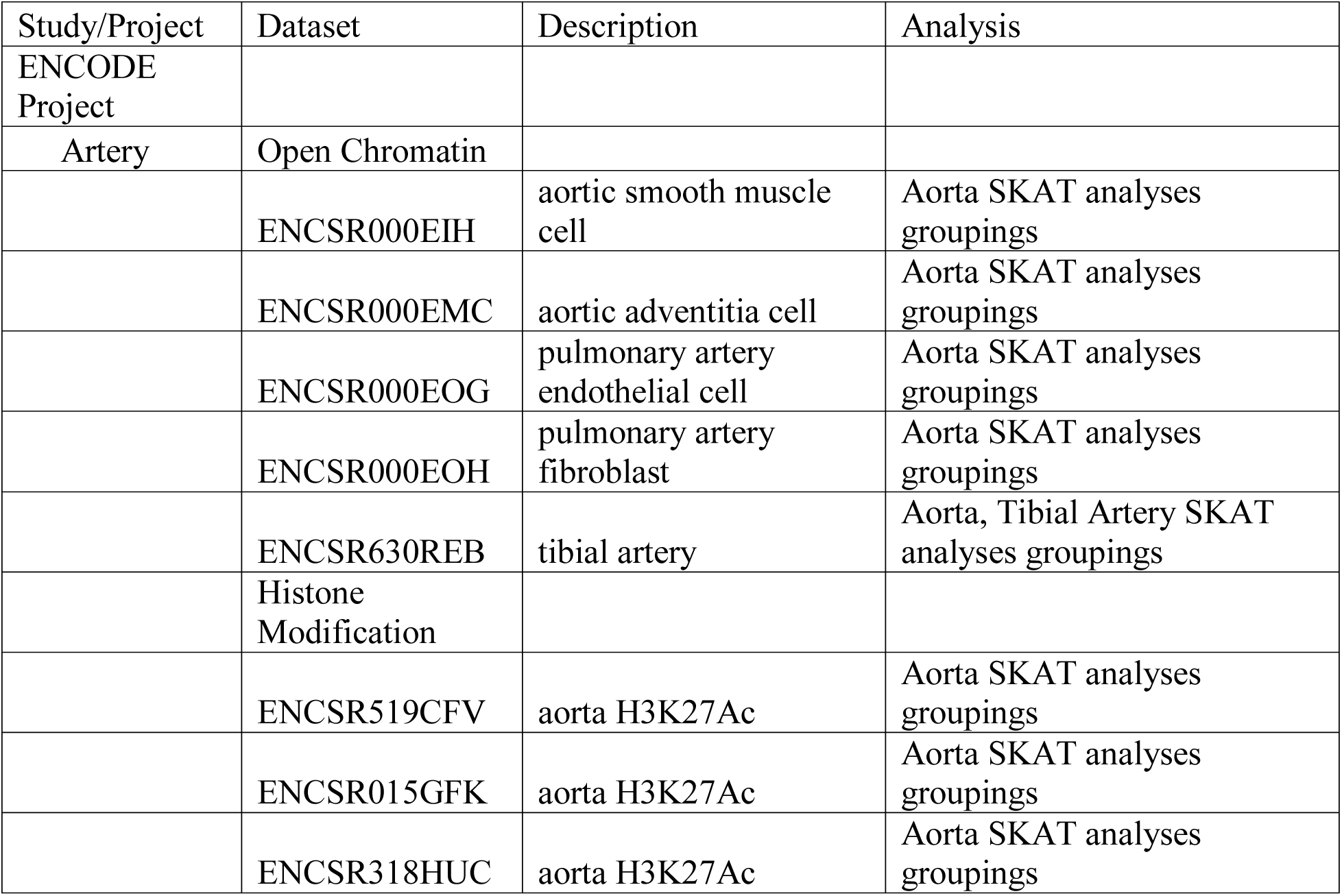
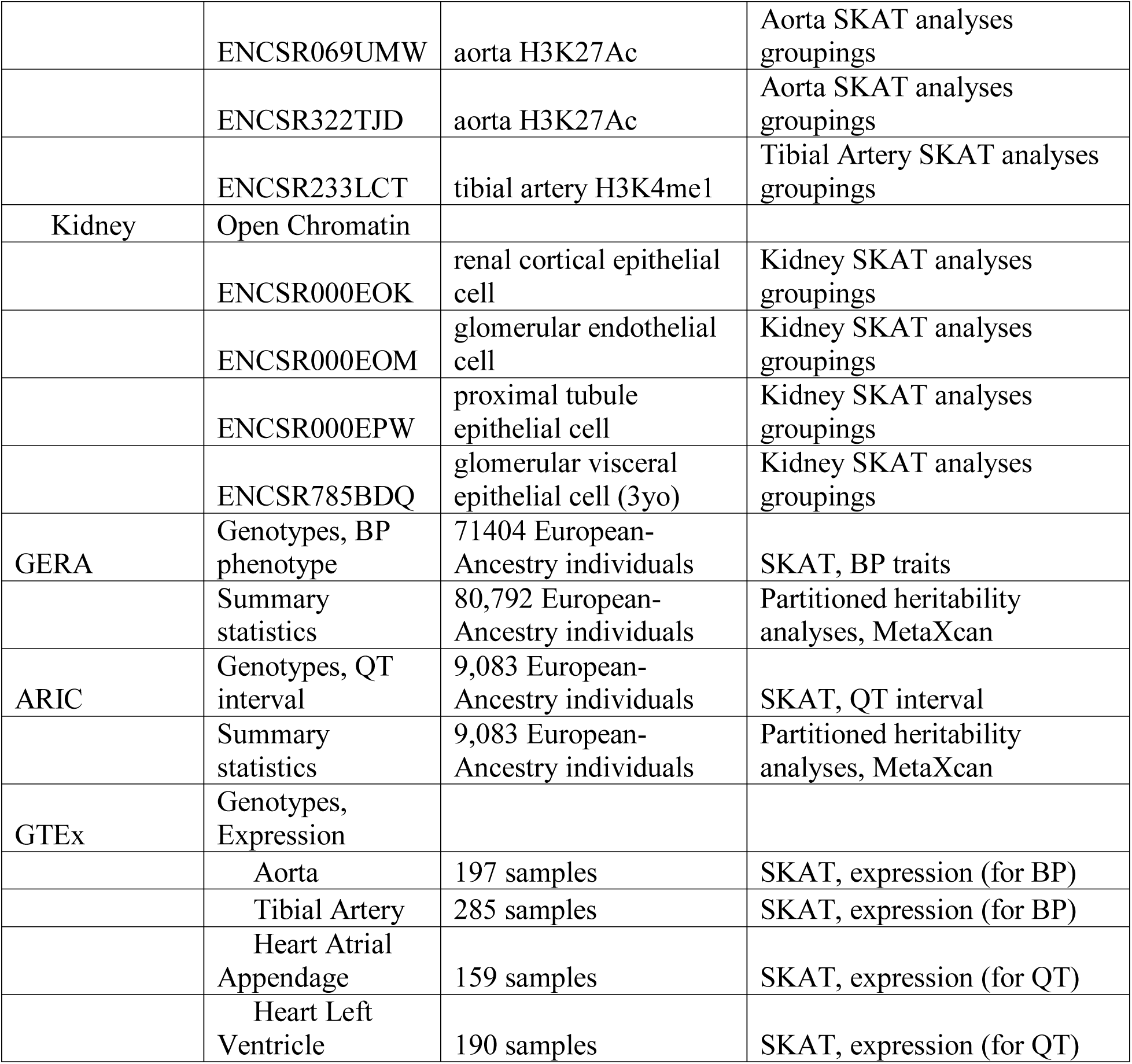
Summary of datasets for analysis in this study.

| Study/Project | Dataset | Description | Analysis |
| --- | --- | --- | --- |
| ENCODE Project |  |  |  |
| Artery | Open Chromatin |  |  |
|  | ENCSR000EIH | aortic smooth muscle cell | Aorta SKAT analyses groupings |
|  | ENCSR000EMC | aortic adventitia cell | Aorta SKAT analyses groupings |
|  | ENCSR000EOG | pulmonary artery endothelial cell | Aorta SKAT analyses groupings |
|  | ENCSR000EOH | pulmonary artery fibroblast | Aorta SKAT analyses groupings |
|  | ENCSR630REB | tibial artery | Aorta, Tibial Artery SKAT analyses groupings |
|  | Histone Modification |  |  |
|  | ENCSR519CFV | aorta H3K27Ac | Aorta SKAT analyses groupings |
|  | ENCSR015GFK | aorta H3K27Ac | Aorta SKAT analyses groupings |
|  | ENCSR318HUC | aorta H3K27Ac | Aorta SKAT analyses groupings |
|  | ENCSR069UMW | aorta H3K27Ac | Aorta SKAT analyses groupings |
|  | ENCSR322TJD | aorta H3K27Ac | Aorta SKAT analyses groupings |
|  | ENCSR233LCT | tibial artery H3K4me1 | Tibial Artery SKAT analyses groupings |
| Kidney | Open Chromatin |  |  |
|  | ENCSR000EOK | renal cortical epithelial cell | Kidney SKAT analyses groupings |
|  | ENCSR000EOM | glomerular endothelial cell | Kidney SKAT analyses groupings |
|  | ENCSR000EPW | proximal tubule epithelial cell | Kidney SKAT analyses groupings |
|  | ENCSR785BDQ | glomerular visceral epithelial cell (3yo) | Kidney SKAT analyses groupings |
| GERA | Genotypes, BP phenotype | 71404 European-Ancestry individuals | SKAT, BP traits |
|  | Summary statistics | 80,792 European-Ancestry individuals | Partitioned heritability analyses, MetaXcan |
| ARIC | Genotypes, QT interval | 9,083 European-Ancestry individuals | SKAT, QT interval |
|  | Summary statistics | 9,083 European-Ancestry individuals | Partitioned heritability analyses, MetaXcan |
| GTE<sub>x</sub> | Genotypes, Expression |  |  |
|  | Aorta | 197 samples | SKAT, expression (for BP) |
|  | Tibial Artery | 285 samples | SKAT, expression (for BP) |
|  | Heart Atrial Appendage | 159 samples | SKAT, expression (for QT) |
|  | Heart Left Ventricle | 190 samples | SKAT, expression (for QT) |

These CRE maps were completed as an extension of the construction of our recent cardiac CRE map [27]. We specifically focused on identifying putative enhancers for the aorta and tibial arteries (see Methods). We subsequently used these maps for training with the software gkm-SVM [28, 29] in order to generate deltaSVM functional scores for all non-coding variants from the 1000 Genomes European ancestry sample, to be tested for association on a gene-level basis. The performance for each model is available in Table S3 (AUC range: 0.84-0.96), with the best performance in the renal cell types. A possible reason for the improved performance of the renal cell types is that the data were from individual cell types as opposed to a mixture of cell types comprising the arteries. The magnitude of the deltaSVM score for a variant reflects its predicted impact on regulatory functional activity, while its sign reflects the prediction with respect to the reference allele. Therefore, to represent the predicted impact of each variant irrespective of allele, we show the distributions of the absolute values of the deltaSVM scores for the arteries and kidney cell types in Fig 1.

**Fig 1.**
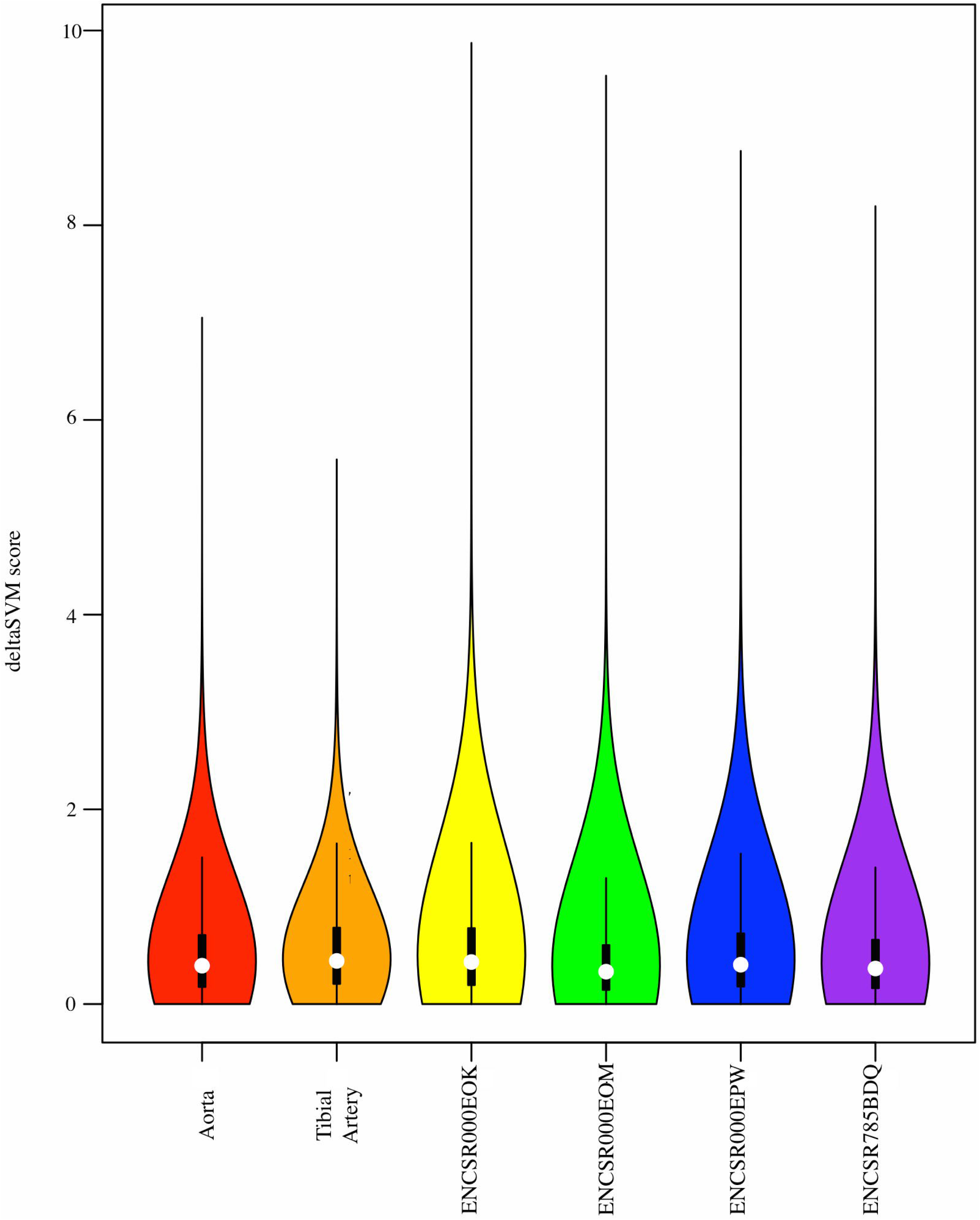
Distributions of deltaSVM scores for each tissue or cell type.

### Tissue-specific gene identification

As our emphasis in this section is to connect a gene’s putative CRE variants to both a phenotype of interest and to its expression in relevant tissues (Fig 2), we first describe the overall analysis scheme as applied to a general phenotype of interest. We then describe how we applied these analyses, first to the QT interval in the ARIC study, as proof of principle to demonstrate the utility of these analyses, and then to our BP traits of interest in the GERA study.

**Fig 2.**
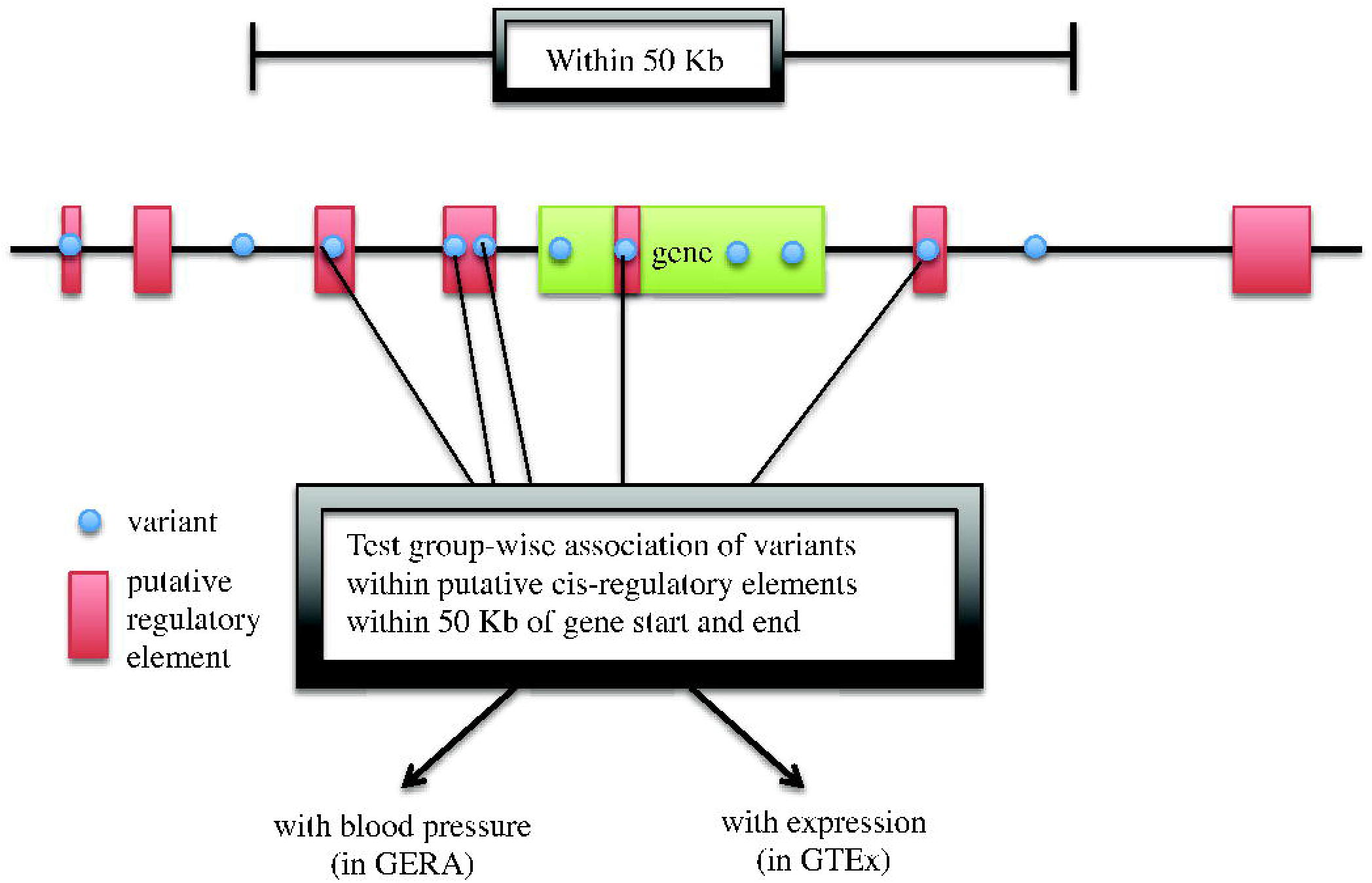
Overview of CRE SKAT analysis.

We defined a gene’s “cis”-regulatory variants in this analysis as those variants falling in putative CREs within 50Kb of the gene’s start and end. We tested their aggregate effect for each gene using SKAT [22], for association tests with the phenotype(s) of interest in the relevant population, SBP and DBP in the GERA study, and QT interval in the ARIC study. SKAT is a test that has generally been used to study groups of variants together and is useful when variants can have bidirectional effects; rare variants are more highly weighted than common variants by default. In addition to the default weights, we ran the analysis using equal weights for all variants. We finally used the tissue- or cell-type-specific deltaSVM scores for the analyzed variants as weights for a customized SKAT test; the score scaling with the effect of the variant on functional regulatory activity.

We then tested these groupings with expression data from GTEx v6p in the tissues of interest to link variants in the genes of interest to their gene expression. The groupings tested in the GTEx data with expression were not always identical to the groupings tested in the GERA or ARIC studies because of differences in imputation quality score filtering, missingness of genotypes from genotype probabilities to hard call conversion, and variants present in the reference populations studied. However, this analysis still connects a given gene to its expression and to the phenotype via a highly overlapping set of CRE variants, and was completed this way to test the most complete set of variants available meeting our criteria. In addition to testing putative regulatory variants with gene expression in GTEx, we used the recently developed MetaXcan [23] software to augment SKAT to identify any new associations by this method. This software predicts the association of gene expression with a phenotype, given genotypes for the population of interest based on training from reference genotypes and expression data.

### Analysis of CREs in QT interval

As mentioned earlier, we considered the cardiac trait QT interval first to demonstrate proof of concept for tissue-specific gene identification. The QT interval is the time in ms between the onset of the Q wave and the end of the T wave in the surface 12-lead electrocardiogram [30], which has ∼30% heritability [31–34]. In our recent work, we have demonstrated that a significant proportion of the heritability is explained by predicted cardiac regulatory variants [27]. We analyzed the genes at previously published QT interval GWAS loci to determine whether or not a heart-specific effect could be observed. Two of the genes with major effects in a GWAS and functionally validated in QT interval heritability are *NOS1AP* [34–36] and *SCN5A* [36, 37]. The full results are presented in the Text S1 results, Table S4, and Figs S1 and S2; to summarize here briefly, we aimed to discover if a heart-specific effect could be revealed for each of these two genes. We observed a heart-specificity for *SCN5A*; *NOS1AP* showed signal across all the cell types in the equal-weighted analyses, though considerably attenuated in some of the deltaSVM-weighted non-heart tissues. Considering both sets of effects, certainly variants with detectable signals present in open chromatin regions specific to the relevant tissue/cell types will allow the detection of a tissue-specific signal, as for *SCN5A*. It also appears, however, that gene-level signals may be captured by analyses in which all variants are weighted equally, and when local open chromatin boundaries across tissues/cell types overlap considerably, especially when variants with strong signals are present within these shared regions. In this situation, we will not necessarily be able to differentiate between different tissue/cell types. Weighting with the tissue-specific deltaSVM scores introduces an additional tier of tissue specificity and is based on global open chromatin differences, and is also not expected to be impacted by linkage disequilibrium (LD) in the ways that the other two weighting schemes are, as the generation of the scores are only dependent on sequence context. Finally, using the default weights shows least concordance with the other two sets of results, indicating that for this analysis, rare variants are not driving the signal as compared to common variants. This is as expected, as we prioritized non-coding variation for these analyses, and the rare variants with larger effects expected to make a detectable contribution are more likely to be in the exome.

### Analysis of CREs at GWAS loci for BP regulation

We then applied these analyses to the tissues of interest for BP regulation, namely aorta, tibial artery, and four kidney cell types, in a subset of 71,404 unrelated GERA EUR individuals. We tested 14,548 genes expressed at RPKM ≥ 0.3 in 197 aorta GTEx samples and 13,963 genes expressed at RPKM ≥ 0.3 in 285 tibial artery GTEx samples for the SKAT analyses. We used summary statistics available from 80,792 individuals[38] to maximize the sample size for which the MetaXcan analyses were run, for the aorta and tibial arteries. Results for each of the arteries are presented in Tables 2-5. In some cases, shared variants drive the positive signal for multiple genes at the same locus; expression in the relevant tissue or cell type may pinpoint a specific gene. However, it may be noted that the genes *CERS5*, *COX14*, and *RP4-605O3.4* are all present at the same locus in the arteries (Tables 2-5), but evidence of expression association is present for many of these genes; this may be indicative of proximal variants affecting different genes, or pleiotropy of single variants affecting expression of multiple genes.

**Table 2.**
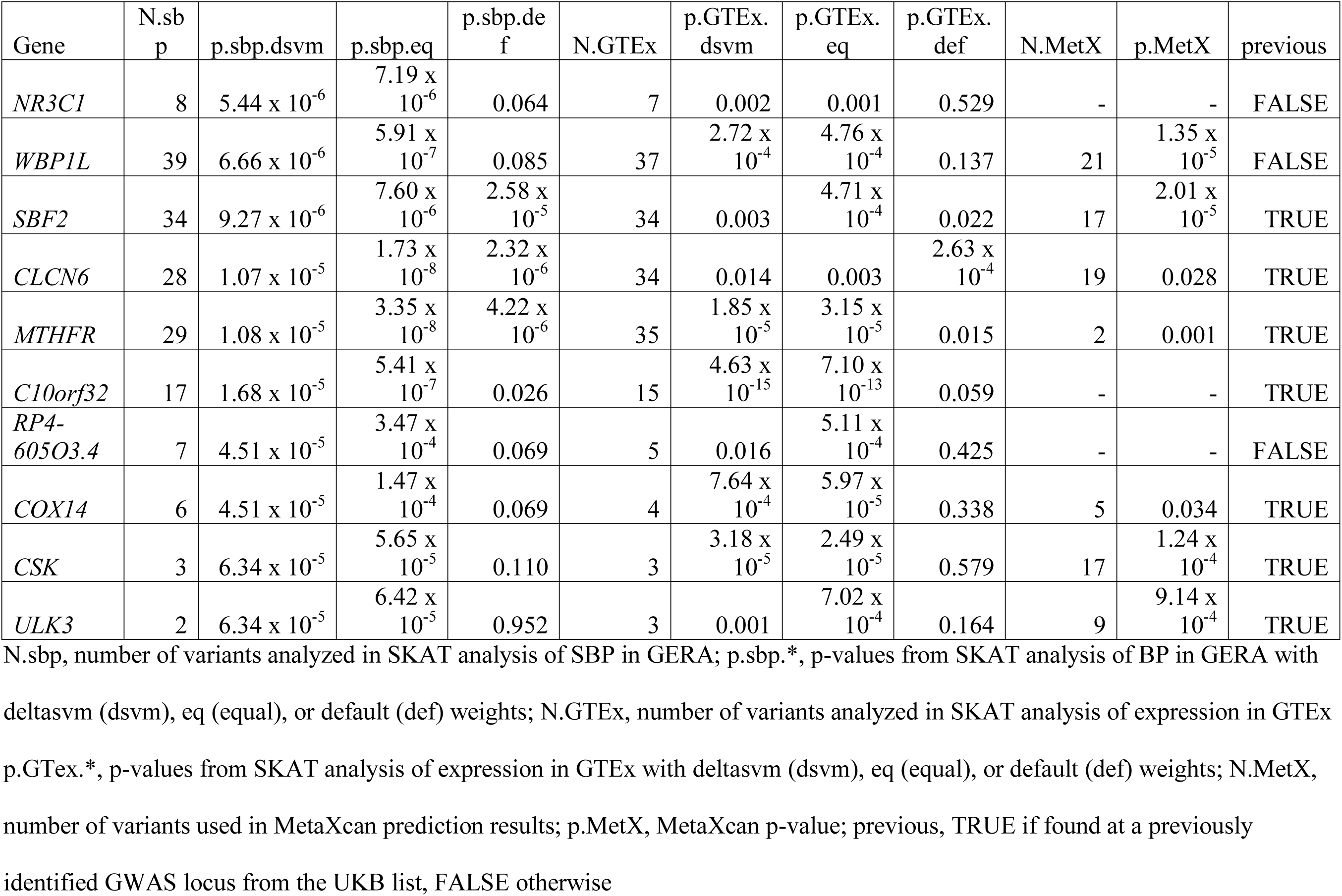
Aorta SBP SKAT and MetaXcan results.

**Table 3.**
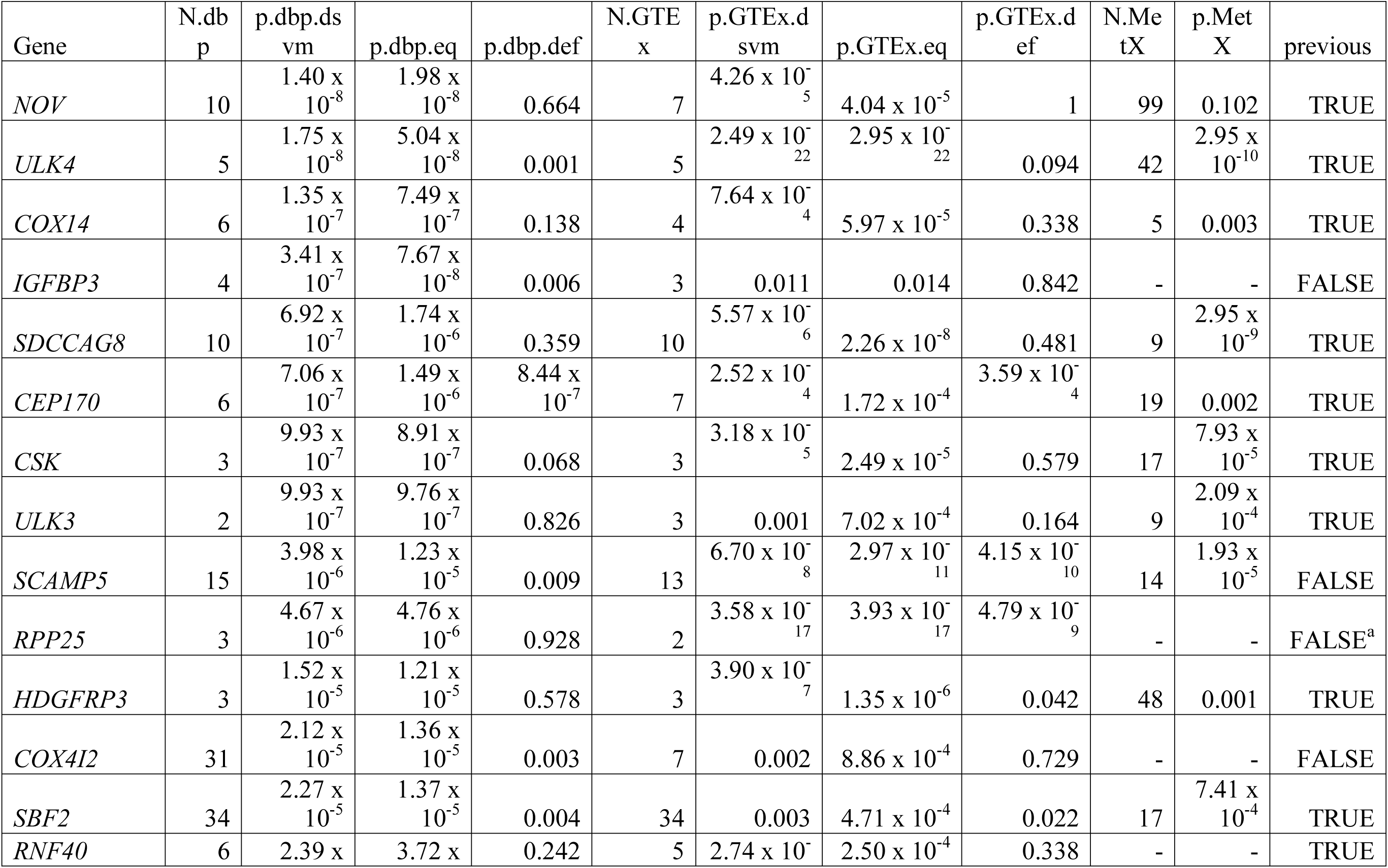
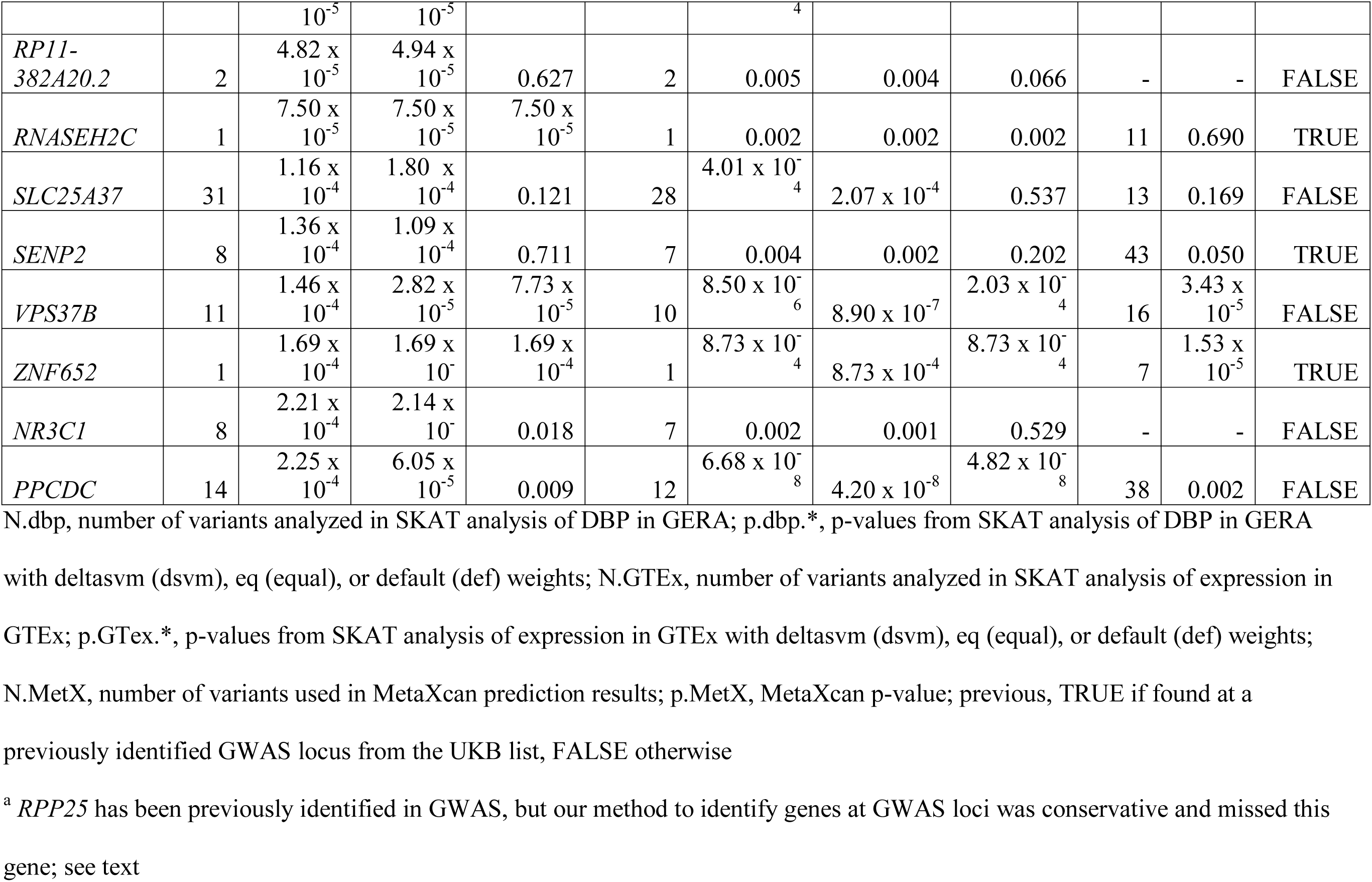
Aorta DBP SKAT and MetaXcan results.

**Table 4.**
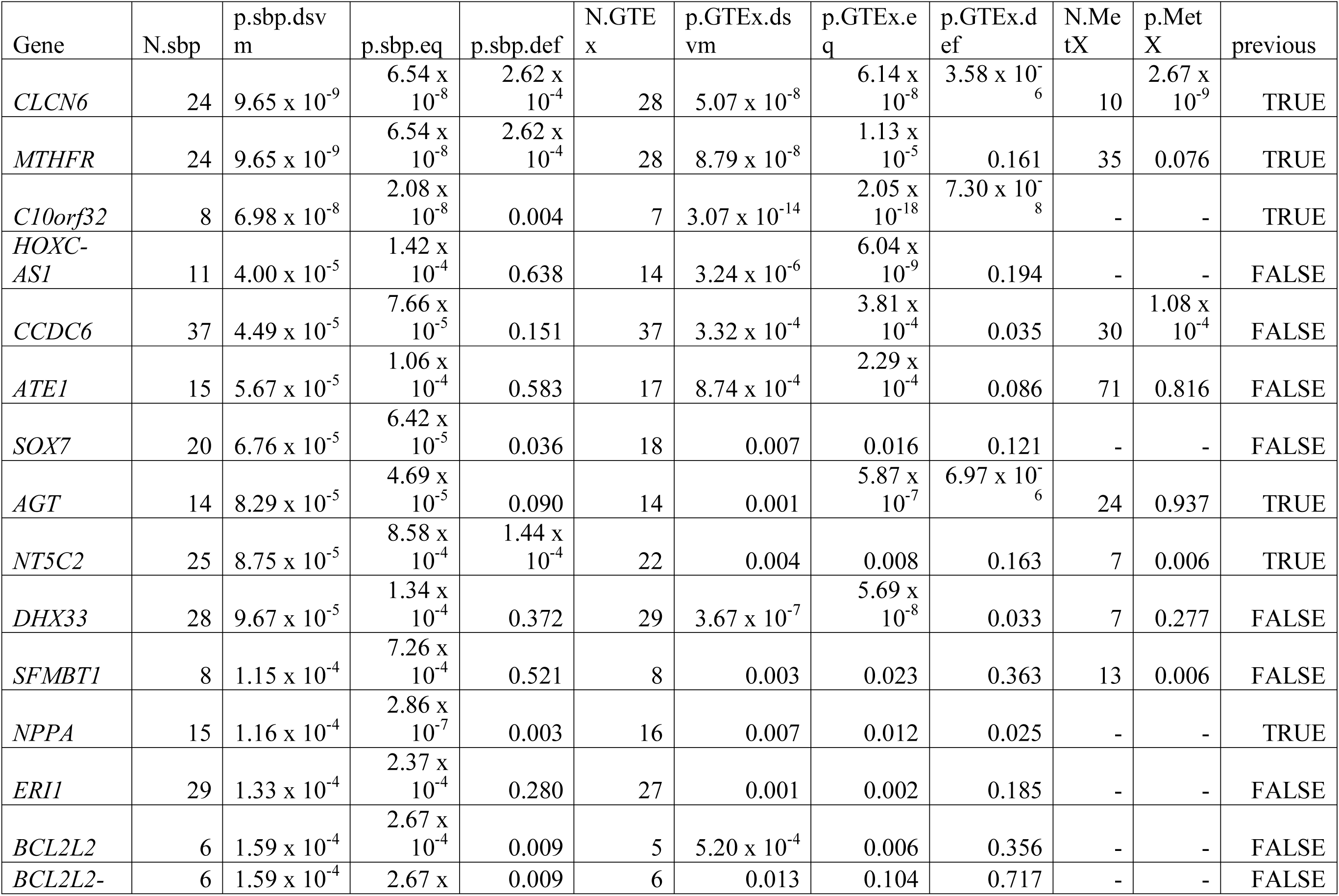
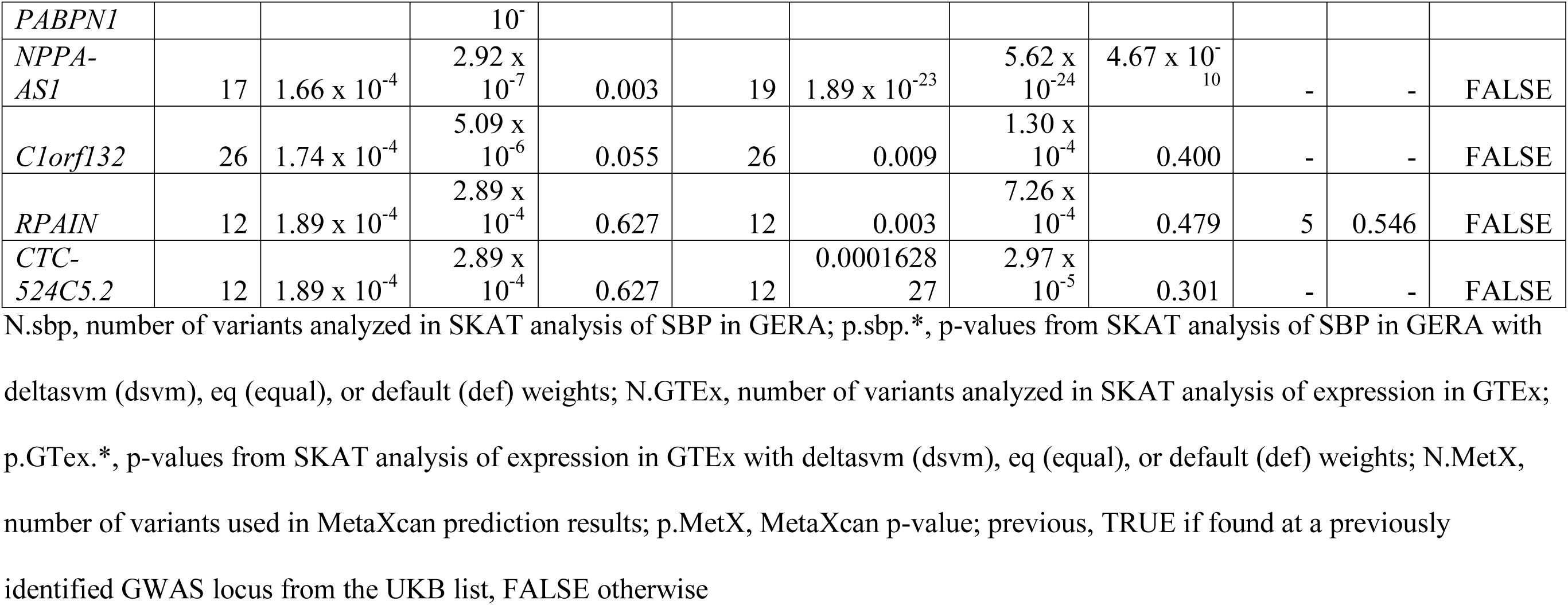
Tibial Artery SBP SKAT and MetaXcan results.

**Table 5.**
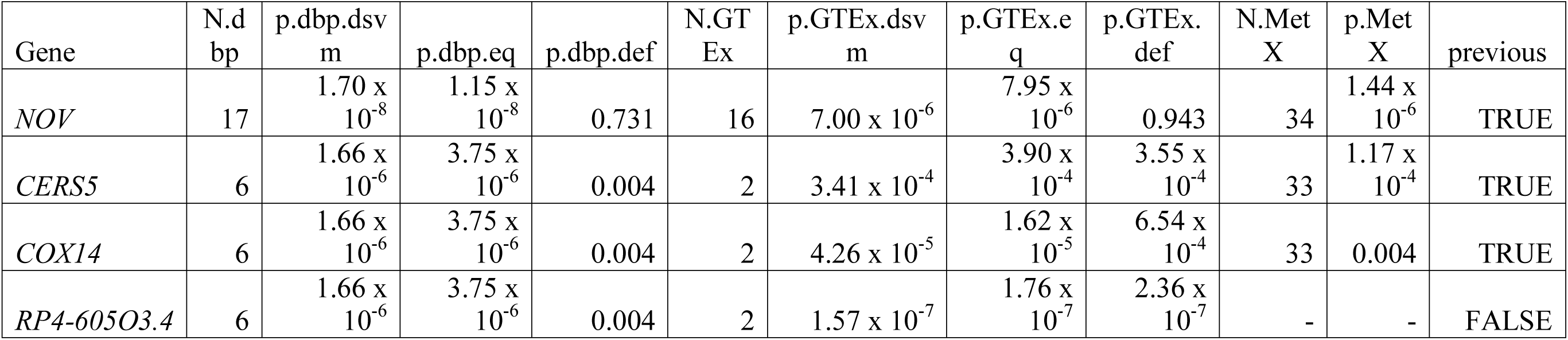
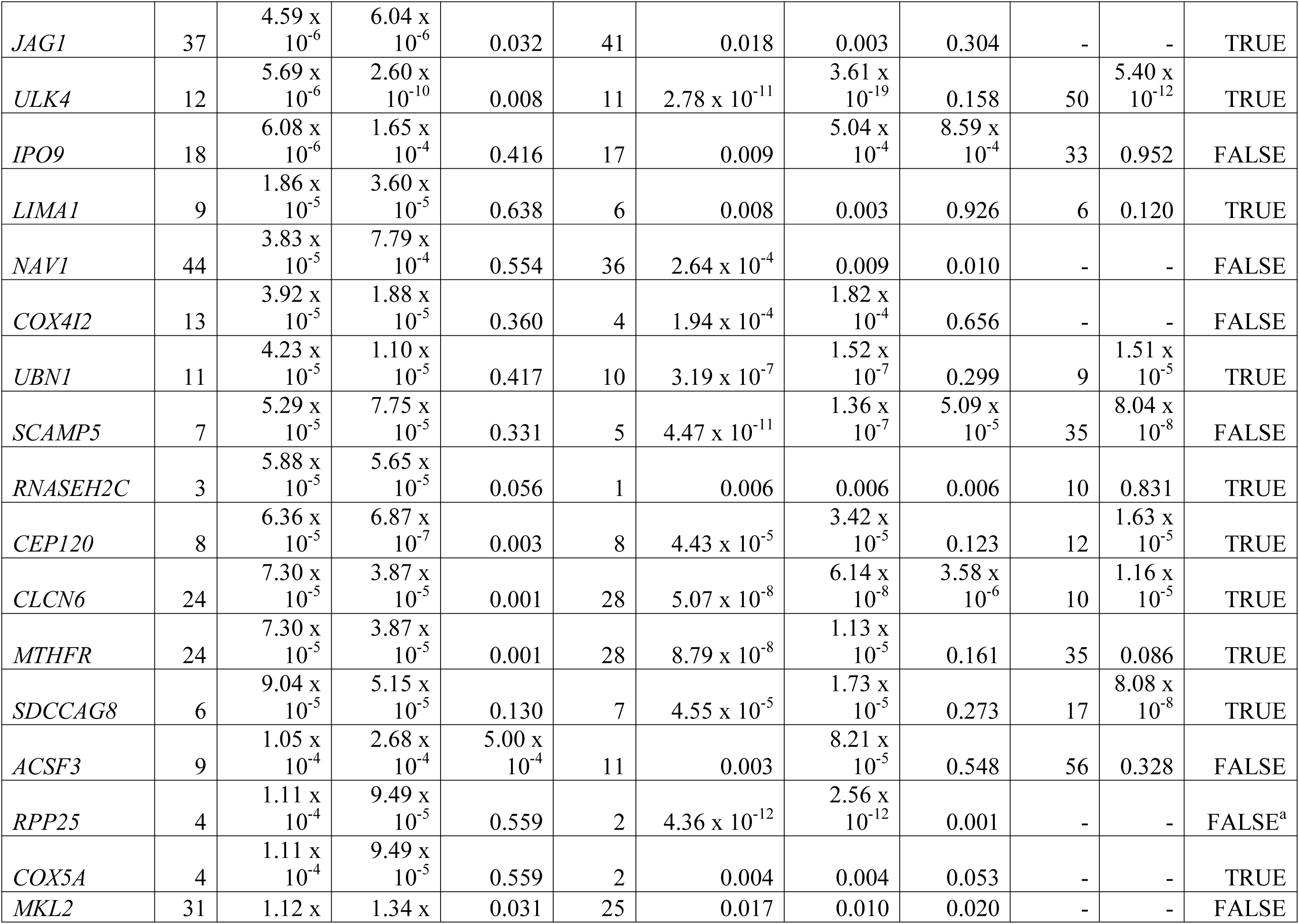
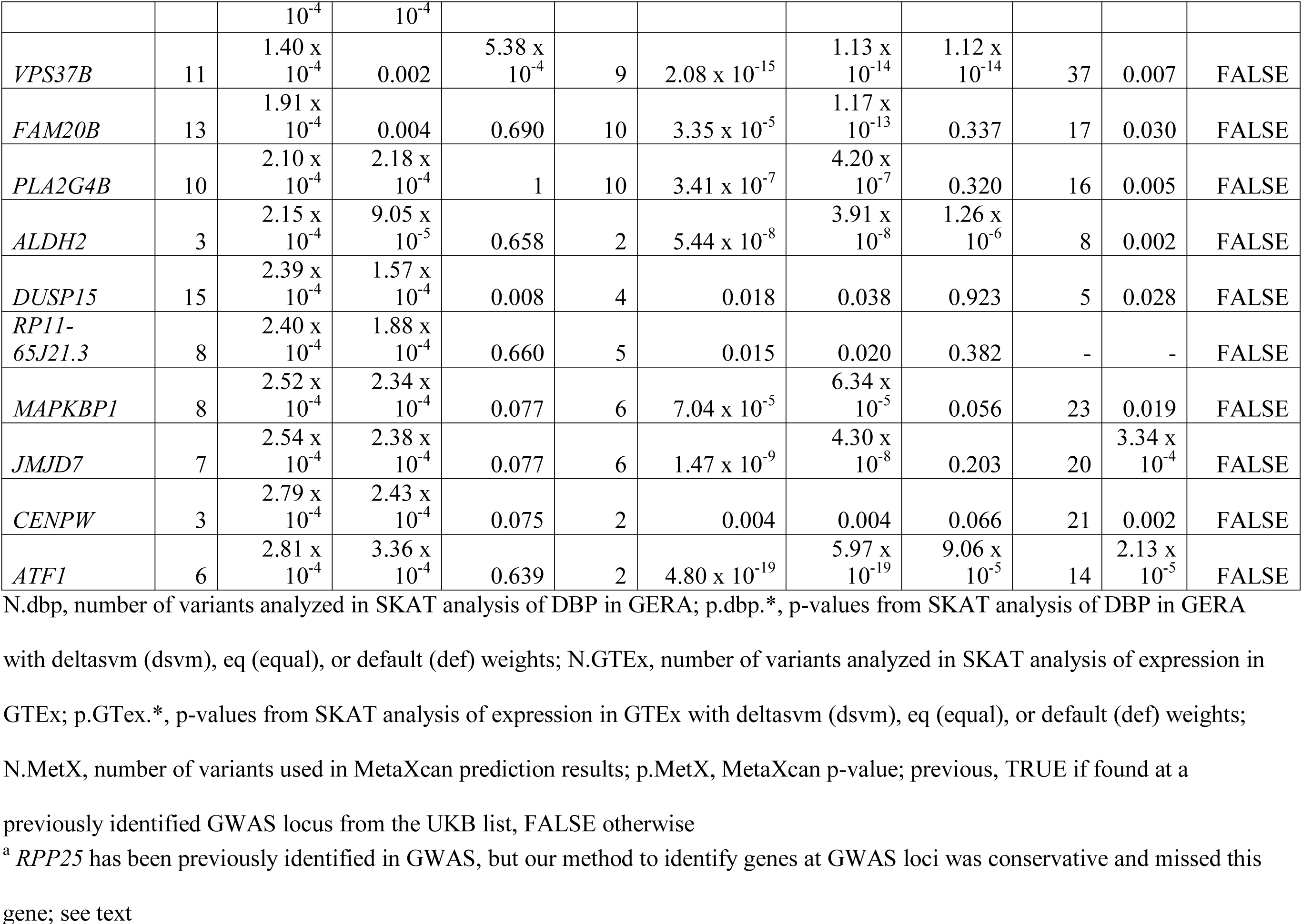
Tibial Artery DBP SKAT and MetaXcan results.

On the whole, the 25 genes reported here across aorta and tibial artery genes have been identified at previous BP loci [39]. While there are several genes in each analysis with interesting associations with BP traits, here we only highlight the genes that have statistical significance of p < 1 x 10^-4^ for both expression and BP in the aorta analyses (Tables 2-3). In our previous work, the aorta was demonstrated as the greatest outlier in an analysis of eQTL enrichment among GTEx tissues for BP traits [38]. These genes include: *MTHFR*[40, 41] (SBP), *C10orf32*[40] (SBP), *CSK* (SBP), *NOV*[42] (DBP), *ULK4*[43] (DBP), *SDCCAG8* (DBP), *SCAMP5* (DBP), *RPP25* (DBP), *HDGFRP3* (DBP), *VPS37B* (DBP), and *PPCDC* (DBP). Most of these genes are present at or near previously replicated BP GWAS loci; *SDCCAG8* was identified as part of Hoffmann et al.[38] It is noteworthy that both *SCAMP5* and *PPCDC* are neighboring genes, but have independent expression support in the same tissue.

### Analysis of CREs at GWAS loci for monogenic BP genes

We also studied the genes involved in monogenic forms of hypotension or hypertension in four kidney cell types available from the ENCODE project (see earlier). As the expression data available for kidney are insufficient, we studied each cell type individually and carried out only SKAT analyses for these genes; the results are in Tables 6-7. The most notable result is that of *CYP17A1*, which shows an effect (p∼10^-5^ – 10^-7^) across all four cell types in the unweighted variants analyses for SBP only, and more specifically, only in the glomerular endothelial cell (ENCSR000EOM) (p.SKAT.dsvm.ENCSR000EOM=3.64 x 10^-8^) in the deltaSVM-weighted results. However, as *C10orf32* is a gene of interest at the same locus, based on the artery results above, we examined and noted that the results are somewhat similar for this gene, although not as striking, due to variant set sharing in the SKAT analyses for these genes (deltaSVM p-values: ENCSR000EOK, 8 variants, p=1.87 x 10^-3^; ENCSR000EOM, 12 variants, p=2.87 x 10^-5^; ENCSR000EPW, 10 variants, p=6.33 x 10^-4^; ENCSR786BDQ, 9 variants, p=0.031). The breakdown of individual variants analyzed for these two genes is in Table S5. The variant rs3824754, with an SBP association p=1.40 x 10^-11^, appears in the groupings of both genes for all four cell types, but has the highest deltaSVM magnitude in the endothelial cell. Additionally, there is a set of four variants with SBP association (p<1 x 10^-4^; rs284853, rs284854, rs284855, rs284856) which only appear in the ENCSR000EOM groupings. We observed that while *CYP17A1* was similarly associated with, or demonstrated evidence of association with, SBP in the deltaSVM and unweighted variants analysis (aorta deltaSVM p=2.06 x 10^-5^, 34 variants; tibial artery deltaSVM p=1.40 x 10^-8^, 15 variants), the analysis of variants in GTEx (33 variants for aorta and 14 variants for tibial artery) did not reflect any significant association (p>0.01). In contrast, *C10orf32* demonstrated association with SBP (aorta deltaSVM p=1.68 x 10^-5^, 17 variants; tibial artery deltaSVM p=6.98 x 10^-8^, 8 variants, Tables 2 and 4) and with expression in GTEx (aorta deltaSVM p=4.63 x 10^-15^, 15 variants; tibial artery deltaSVM p=3.07 x 10^-14^, 7 variants, Tables 2 and 4). The same four variants unique to the ENCSR000EOM groupings above with strong associations with SBP are also present in the artery groupings. Three of these variants (rs284854, rs284855, rs284856) are eQTLs for *C10orf32* in the aorta and tibial arteries; these variants, however, do not show association with *CYP17A1* expression in these tissues (all p > 0.03 for aorta, all p > 0.21 for tibial artery, from eQTL data available from the GTEx portal (https://www.gtexportal.org/), accessed 09/08/17). Additionally, as the *CYP17A1* gene primarily demonstrates an adrenal effect in the monogenic disorder [44], we also examined the associations of these three variants in the GTEx portal with adrenal gland expression data for both genes; all have p > 0.26 for *CYP17A1* and p > 0.04 for *C10orf32*. This may reflect an endothelial-cell-specific effect for *C10orf32* rather than a tissue-type effect, especially as this locus has been identified in several previous BP GWAS studies, [40,43,45–47]; it may also not be very informative for the kidney, though suitable expression data for kidney would be required to assess this.

**Table 6.** Kidney SBP SKAT results.

| Gene | Experiment<sup>a</sup> | N.sbp | p.sbp.dsvm | p.sbp.eq | p.sbp.def |
| --- | --- | --- | --- | --- | --- |
| *BSND* | ENCSR000EOK | 13 | 0.836 | 0.796 | 0.611 |
| *CASR* | ENCSR000EOK | 21 | 0.628 | 0.794 | 0.605 |
| *CLCNKA* | ENCSR000EOK | 8 | 0.444 | 0.502 | 0.426 |
| *CLCNKB* | ENCSR000EOK | 9 | 0.475 | 0.382 | 0.454 |
| *CUL3* | ENCSR000EOK | 18 | 0.318 | 0.388 | 0.049 |
| *CYP11B1* | ENCSR000EOK | 12 | 0.020 | 0.016 | 0.001 |
| *CYP11B2* | ENCSR000EOK | 10 | 0.014 | 0.014 | 0.003 |
| *CYP17A1* | ENCSR000EOK | 7 | 0.069 | $1.11 \times 10^{-6}$ | 0.018 |
| *HSD11B2* | ENCSR000EOK | 4 | 0.150 | 0.104 | 0.248 |
| *KCNJ1* | ENCSR000EOK | 13 | 0.992 | 0.958 | 0.789 |
| *KCNJ5* | ENCSR000EOK | 11 | 0.550 | 0.742 | 0.982 |
| *KLHL3* | ENCSR000EOK | 12 | 0.573 | 0.431 | 0.345 |
| *NR3C2* | ENCSR000EOK | 26 | 0.375 | 0.270 | 0.148 |
| *SCNN1A* | ENCSR000EOK | 7 | 0.502 | 0.452 | 0.767 |
| *SCNN1B* | ENCSR000EOK | 7 | 0.053 | 0.310 | 0.479 |
| *SCNN1G* | ENCSR000EOK | 8 | 0.170 | 0.276 | 1.000 |
| *SLC12A1* | ENCSR000EOK | 6 | 1.000 | 0.909 | 0.713 |
| *SLC12A3* | ENCSR000EOK | 16 | 0.233 | 0.168 | 0.491 |
| *WNK1* | ENCSR000EOK | 11 | 0.365 | 0.557 | 0.637 |
| *WNK4* | ENCSR000EOK | 1 | 0.010 | 0.010 | 0.010 |
| *BSND* | ENCSR000EOM | 12 | 0.861 | 0.737 | 0.562 |
| *CASR* | ENCSR000EOM | 2 | 0.860 | 0.790 | 0.672 |
| *CLCNKA* | ENCSR000EOM | 10 | 0.688 | 0.657 | 0.540 |
| *CLCNKB* | ENCSR000EOM | 10 | 0.688 | 0.657 | 0.540 |
| *CUL3* | ENCSR000EOM | 8 | 0.081 | 0.250 | 0.044 |
| *CYP11B1* | ENCSR000EOM | 4 | 0.018 | 0.018 | 0.281 |
| *CYP11B2* | ENCSR000EOM | 3 | 0.015 | 0.015 | 0.281 |
| *CYP17A1* | ENCSR000EOM | 11 | $3.64 \times 10^{-8}$ | $2.88 \times 10^{-7}$ | 0.013 |
| *HSD11B2* | ENCSR000EOM | 6 | 0.024 | 0.044 | 0.595 |
| *KCNJ1* | ENCSR000EOM | 9 | 0.086 | 0.554 | 0.060 |
| *KCNJ5* | ENCSR000EOM | 6 | 0.084 | 0.150 | 0.163 |
| *KLHL3* | ENCSR000EOM | 12 | 0.065 | 0.012 | 0.208 |
| *NR3C2* | ENCSR000EOM | 10 | 0.278 | 0.126 | 0.149 |
| *SCNN1A* | ENCSR000EOM | 5 | 0.517 | 0.572 | 0.727 |
| *SCNN1B* | ENCSR000EOM | 5 | 0.815 | 0.930 | 0.873 |
| *SCNN1G* | ENCSR000EOM | 5 | 0.353 | 0.298 | 0.540 |
| *SLC12A1* | ENCSR000EOM | 4 | 0.742 | 0.896 | 0.704 |
| *SLC12A3* | ENCSR000EOM | 12 | 0.062 | 0.077 | 0.245 |
| *WNK1* | ENCSR000EOM | 7 | 0.151 | 0.158 | 0.864 |
| *WNK4* | ENCSR000EOM | 1 | 0.010 | 0.010 | 0.010 |
| *BSND* | ENCSR000EPW | 10 | 0.649 | 0.702 | 0.591 |
| *CASR* | ENCSR000EPW | 11 | 0.665 | 0.798 | 0.335 |
| *CLCNKA* | ENCSR000EPW | 3 | 0.190 | 0.203 | 0.433 |
| *CLCNKB* | ENCSR000EPW | 4 | 0.268 | 0.176 | 0.464 |
| *CUL3* | ENCSR000EPW | 8 | 0.030 | 0.195 | 0.039 |
| *CYP11B1* | ENCSR000EPW | 3 | 0.015 | 0.015 | 0.281 |
| *CYP11B2* | ENCSR000EPW | 5 | 0.015 | 0.017 | 0.488 |
| *CYP17A1* | ENCSR000EPW | 9 | 0.006 | $9.92 \times 10^{-7}$ | 0.045 |
| *HSD11B2* | ENCSR000EPW | 1 | 0.051 | 0.051 | 0.051 |
| *KCNJ1* | ENCSR000EPW | 7 | 0.960 | 0.954 | 0.411 |
| *KCNJ5* | ENCSR000EPW | 10 | 0.519 | 0.699 | 0.417 |
| *KLHL3* | ENCSR000EPW | 8 | 0.380 | 0.525 | 0.428 |
| *NR3C2* | ENCSR000EPW | 24 | 0.264 | 0.189 | 0.027 |
| *SCNN1A* | ENCSR000EPW | 6 | 0.492 | 0.360 | 0.767 |
| *SCNN1B* | ENCSR000EPW | 7 | 0.141 | 0.181 | 0.482 |
| *SCNN1G* | ENCSR000EPW | 6 | 0.778 | 0.565 | 1.000 |
| *SLC12A1* | ENCSR000EPW | 4 | 1.000 | 0.896 | 0.704 |
| *SLC12A3* | ENCSR000EPW | 13 | 0.135 | 0.116 | 0.358 |
| *WNK1* | ENCSR000EPW | 11 | 0.526 | 0.515 | 0.054 |
| *WNK4* | ENCSR000EPW | 1 | 0.010 | 0.010 | 0.010 |
| *BSND* | ENCSR785BDQ | 10 | 0.888 | 0.854 | 0.668 |
| *CASR* | ENCSR785BDQ | 1 | 0.465 | 0.465 | 0.465 |
| *CLCNKA* | ENCSR785BDQ | 10 | 0.466 | 0.606 | 0.682 |
| *CLCNKB* | ENCSR785BDQ | 11 | 0.457 | 0.476 | 0.695 |
| *CUL3* | ENCSR785BDQ | 2 | 0.641 | 0.314 | 0.982 |
| *CYP11B1* | ENCSR785BDQ | 4 | 0.387 | 0.353 | 0.026 |
| *CYP11B2* | ENCSR785BDQ | 2 | 0.025 | 0.026 | 0.996 |
| *CYP17A1* | ENCSR785BDQ | 12 | 0.008 | $3.93 \times 10^{-5}$ | 0.020 |
| *HSD11B2* | ENCSR785BDQ | 6 | 0.205 | 0.067 | 0.061 |
| *KCNJ1* | ENCSR785BDQ | 7 | 0.961 | 0.879 | 0.400 |
| *KCNJ5* | ENCSR785BDQ | 2 | 1.000 | 1.000 | 1.000 |
| *KLHL3* | ENCSR785BDQ | 10 | 0.498 | 0.126 | 0.888 |
| *NR3C2* | ENCSR785BDQ | 9 | 0.314 | 0.290 | 0.037 |
| *SCNN1A* | ENCSR785BDQ | 9 | 0.521 | 0.700 | 0.980 |
| *SCNN1B* | ENCSR785BDQ | 4 | 1.000 | 1.000 | 0.983 |
| *SCNN1G* | ENCSR785BDQ | 6 | 0.727 | 0.565 | 1.000 |
| *SLC12A1* | ENCSR785BDQ | 2 | 0.848 | 0.848 | 0.848 |
| *SLC12A3* | ENCSR785BDQ | 28 | 0.250 | 0.440 | 0.666 |
| *WNK1* | ENCSR785BDQ | 7 | 0.289 | 0.414 | 0.128 |
| *WNK4* | ENCSR785BDQ | - | - | - | - |

N.sbp, number of variants analyzed in SKAT analysis of SBP in GERA; p.sbp.\*, p-values from
SKAT analysis of BP in GERA with deltasvm (dsvm), eq (equal), or default (def) weights
<sup>a</sup> Experiments: ENCSR000EOK, renal cortical epithelial cell; ENCSR000EOM, glomerular
endothelial cell; ENCSR000EPW, epithelial cell of proximal tubule; ENCSR785BDQ,
glomerular visceral epithelial cell

**Table 7.** Kidney DBP SKAT results.

| Gene | Experiment<sup>a</sup> | N.dbp | p.dbp.dsvm | p.dbp.eq | p.dbp.def |
| --- | --- | --- | --- | --- | --- |
| *BSND* | ENCSR000EOK | 13 | 0.938 | 0.831 | 0.061 |
| *CASR* | ENCSR000EOK | 21 | 0.990 | 0.986 | 0.856 |
| *CLCNKA* | ENCSR000EOK | 8 | 0.126 | 0.283 | 0.108 |
| *CLCNKB* | ENCSR000EOK | 9 | 0.131 | 0.410 | 0.137 |
| *CUL3* | ENCSR000EOK | 18 | 0.270 | 0.183 | 0.855 |
| *CYP11B1* | ENCSR000EOK | 12 | 0.102 | 0.095 | 0.013 |
| *CYP11B2* | ENCSR000EOK | 10 | 0.091 | 0.094 | 0.023 |
| *CYP17A1* | ENCSR000EOK | 7 | 0.505 | 0.189 | 0.337 |
| *HSD11B2* | ENCSR000EOK | 4 | 0.045 | 0.023 | 0.114 |
| *KCNJ1* | ENCSR000EOK | 13 | 0.969 | 0.879 | 0.398 |
| *KCNJ5* | ENCSR000EOK | 11 | 0.722 | 0.598 | 0.487 |
| *KLHL3* | ENCSR000EOK | 12 | 0.835 | 0.922 | 0.622 |
| *NR3C2* | ENCSR000EOK | 26 | 0.730 | 0.766 | 0.104 |
| *SCNN1A* | ENCSR000EOK | 7 | 0.751 | 0.621 | 0.599 |
| *SCNN1B* | ENCSR000EOK | 7 | 3.33 x 10<sup>-4</sup> | 0.012 | 0.053 |
| *SCNN1G* | ENCSR000EOK | 8 | 0.074 | 0.027 | 0.461 |
| *SLC12A1* | ENCSR000EOK | 6 | 0.361 | 0.259 | 0.860 |
| *SLC12A3* | ENCSR000EOK | 16 | 0.069 | 0.072 | 0.175 |
| *WNK1* | ENCSR000EOK | 11 | 0.375 | 0.300 | 0.442 |
| *WNK4* | ENCSR000EOK | 1 | 0.828 | 0.828 | 0.828 |
| *BSND* | ENCSR000EOM | 12 | 0.613 | 0.776 | 0.052 |
| *CASR* | ENCSR000EOM | 2 | 0.877 | 0.825 | 0.759 |
| *CLCNKA* | ENCSR000EOM | 10 | 0.349 | 0.546 | 0.133 |
| *CLCNKB* | ENCSR000EOM | 10 | 0.349 | 0.546 | 0.133 |
| *CUL3* | ENCSR000EOM | 8 | 0.290 | 0.266 | 0.798 |
| *CYP11B1* | ENCSR000EOM | 4 | 0.117 | 0.129 | 0.689 |
| *CYP11B2* | ENCSR000EOM | 3 | 0.096 | 0.105 | 0.689 |
| *CYP17A1* | ENCSR000EOM | 11 | 0.053 | 0.076 | 0.187 |
| *HSD11B2* | ENCSR000EOM | 6 | 0.012 | 0.012 | 0.387 |
| *KCNJ1* | ENCSR000EOM | 9 | 0.098 | 0.317 | 0.101 |
| *KCNJ5* | ENCSR000EOM | 6 | 0.106 | 0.499 | 0.598 |
| *KLHL3* | ENCSR000EOM | 12 | 0.897 | 0.924 | 0.515 |
| *NR3C2* | ENCSR000EOM | 10 | 0.671 | 0.302 | 0.072 |
| *SCNN1A* | ENCSR000EOM | 5 | 0.782 | 0.709 | 0.547 |
| *SCNN1B* | ENCSR000EOM | 5 | 0.763 | 0.907 | 0.913 |
| *SCNN1G* | ENCSR000EOM | 5 | 0.167 | 0.085 | 0.920 |
| *SLC12A1* | ENCSR000EOM | 4 | 0.118 | 0.244 | 0.865 |
| *SLC12A3* | ENCSR000EOM | 12 | 0.008 | 0.044 | 0.234 |
| *WNK1* | ENCSR000EOM | 7 | 0.499 | 0.589 | 1.000 |
| *WNK4* | ENCSR000EOM | 1 | 0.828 | 0.828 | 0.828 |
| *BSND* | ENCSR000EPW | 10 | 0.866 | 0.769 | 0.062 |
| *CASR* | ENCSR000EPW | 11 | 0.921 | 0.845 | 0.872 |
| *CLCNKA* | ENCSR000EPW | 3 | 0.431 | 0.438 | 0.367 |
| *CLCNKB* | ENCSR000EPW | 4 | 0.712 | 0.639 | 0.413 |
| *CUL3* | ENCSR000EPW | 8 | 0.239 | 0.085 | 0.783 |
| *CYP11B1* | ENCSR000EPW | 3 | 0.104 | 0.105 | 0.689 |
| *CYP11B2* | ENCSR000EPW | 5 | 0.104 | 0.102 | 0.599 |
| *CYP17A1* | ENCSR000EPW | 9 | 0.360 | 0.185 | 0.262 |
| *HSD11B2* | ENCSR000EPW | 1 | 0.006 | 0.006 | 0.006 |
| *KCNJ1* | ENCSR000EPW | 7 | 0.709 | 0.873 | 0.366 |
| *KCNJ5* | ENCSR000EPW | 10 | 0.680 | 0.546 | 0.190 |
| *KLHL3* | ENCSR000EPW | 8 | 0.817 | 0.867 | 0.448 |
| *NR3C2* | ENCSR000EPW | 24 | 0.489 | 0.374 | 0.032 |
| *SCNN1A* | ENCSR000EPW | 6 | 0.733 | 0.446 | 0.599 |
| *SCNN1B* | ENCSR000EPW | 7 | 0.079 | 0.017 | 0.053 |
| *SCNN1G* | ENCSR000EPW | 6 | 0.647 | 0.061 | 0.297 |
| *SLC12A1* | ENCSR000EPW | 4 | 0.415 | 0.244 | 0.865 |
| *SLC12A3* | ENCSR000EPW | 13 | 0.017 | 0.050 | 0.217 |
| *WNK1* | ENCSR000EPW | 11 | 0.300 | 0.173 | 0.031 |
| *WNK4* | ENCSR000EPW | 1 | 0.828 | 0.828 | 0.828 |
| *BSND* | ENCSR785BDQ | 10 | 0.645 | 0.492 | 0.062 |
| *CASR* | ENCSR785BDQ | 1 | 0.967 | 0.967 | 0.967 |
| *CLCNKA* | ENCSR785BDQ | 10 | 0.291 | 0.457 | 0.268 |
| *CLCNKB* | ENCSR785BDQ | 11 | 0.296 | 0.568 | 0.303 |
| *CUL3* | ENCSR785BDQ | 2 | 0.145 | 0.040 | 0.248 |
| *CYP11B1* | ENCSR785BDQ | 4 | 0.154 | 0.134 | 0.003 |
| *CYP11B2* | ENCSR785BDQ | 2 | 0.105 | 0.107 | 0.508 |
| *CYP17A1* | ENCSR785BDQ | 12 | 0.551 | 0.218 | 0.355 |
| *HSD11B2* | ENCSR785BDQ | 6 | 0.079 | 0.008 | 0.003 |
| *KCNJ1* | ENCSR785BDQ | 7 | 0.848 | 0.795 | 0.128 |
| *KCNJ5* | ENCSR785BDQ | 2 | 1.000 | 1.000 | 0.875 |
| *KLHL3* | ENCSR785BDQ | 10 | 0.802 | 0.778 | 0.424 |
| *NR3C2* | ENCSR785BDQ | 9 | 0.877 | 0.784 | 0.019 |
| *SCNN1A* | ENCSR785BDQ | 9 | 0.789 | 0.949 | 0.925 |
| *SCNN1B* | ENCSR785BDQ | 4 | 0.345 | 0.445 | 0.555 |
| *SCNN1G* | ENCSR785BDQ | 6 | 0.083 | 0.061 | 0.297 |
| *SLC12A1* | ENCSR785BDQ | 2 | 0.282 | 0.282 | 0.282 |
| *SLC12A3* | ENCSR785BDQ | 28 | 0.044 | 0.415 | 0.393 |
| *WNK1* | ENCSR785BDQ | 7 | 0.741 | 0.416 | 0.079 |
| *WNK4* | ENCSR785BDQ | - | - | - | - |

N.dbp, number of variants analyzed in SKAT analysis of DBP in GERA; p.dbp.\*, p-values from
SKAT analysis of DBP in GERA with deltasvm (dsvm), eq (equal), or default (def) weights
<sup>a</sup> Experiments: ENCSR000EOK, renal cortical epithelial cell; ENCSR000EOM, glomerular
endothelial cell; ENCSR000EPW, epithelial cell of proximal tubule; ENCSR785BDQ,
glomerular visceral epithelial cell

Quantile-quantile (QQ) plots are shown in Figs S3-S14 for each of the six tissues and two BP phenotypes, and each of three delta SVM weighting schemes. Although the dsvm weighting scheme demonstrates a greater enrichment of genes than the default weighting scheme, the equal weighting scheme marginally presents the greatest enrichment. In many cases, deltaSVM discriminates between different tissue/cell types while equal-weighted results do not; this is especially clear with the QT interval results.

We analyzed the union of genes that met significance in the association analysis with BP, regardless of association of expression, to maximize our gene list for annotation, using DAVID 6.8 [48, 49]. The results are shown in Table S6.

## Discussion

Our previous genetic analyses identified the aorta and tibial arteries as relevant to blood pressure regulation [38]. In this study we have now identified several genes with regulatory variants linking significantly to both BP traits and to expression data in these tissues, most at previously replicated BP loci. Although the involvement of the kidney is well established in BP regulation through physiological evidence, we sought to identify genes at any of the hundreds of BP GWAS loci in a broader set of tissues. We examined groupings of multiple proximal and putatively causal variants defined around genes within a single tissue in order to identify specific genes of interest. We also examined QT interval genes at previous GWAS loci to highlight the identification of functionally characterized genes for this trait.

We identified several genes of potential interest to the aorta/arteries for BP, mostly at previously identified GWAS loci: *MTHFR*, *C10orf32*, *CSK*, *NOV*, *ULK4*, *SDCCAG8*, *SCAMP5*, *RPP25*, *HDGFRP3*, *VPS37B*, and *PPCDC*. We note here that our method of identifying genes at previous loci was conservative: *RPP25* was not present in this list, but is present just outside the TAD boundary used. In addition to its role in the progression of various cancers, the *NOV* gene has been identified as a player in angiogenesis [50, 51] and vascular homeostasis [52]. The *ULK4* gene has been previously associated with DBP [43], and variation in this gene has also been associated with aortic disease and acute aortic dissections [53]. The association of a homozygous variant (C677T) in its neighboring gene, *MTHFR*, has long been associated with BP and vascular disease [54–57]; more generally, this locus has been identified in large BP GWAS [40, 41]. The locus including *C10orf32* has been identified previously [40] and neighbors the well-studied *CYP17A1* gene. Though we initially examined only the latter among kidney cell types, because of its known role in monogenic hypertension, we note that both genes show BP association in endothelial contexts as well, but it is *C10orf32* that has strong expression support in the artery datasets in our study, while *CYP17A1* does not [58]. The gene *SDCCAG8* is a centrosomal protein linked with nephronophthisis-related ciliopathies (OMIM: Senior-Loken Syndrome 7, 613615; Bardet-Biedl syndrome-16, 615993, and Airik et al. [59]), and is expressed in the kidney and lung epithelia [59]. The *CSK* gene, encoding a tyrosine kinase, is at a previous BP GWAS locus [60] and has been found to be associated with SBP in young children [61]; there is also prior evidence through experiments in mouse aortas that this gene regulates blood pressure through Src [62]. Finally, the *SCAMP5* and *PPCDC* genes (within the same locus) [40], and *RPP25* [43], are previously identified BP genes.

As mentioned above, one major limitation in our study is the statistical power of the SKAT eQTL analysis, with small sample sizes available for each of the GTEx tissues. The power of implicating effects for a given tissue also depends on its total contribution, and the numbers of eQTLs identified. The requirement in our study that a gene meet significance for both BP and expression therefore produced a more conservative list. However, the QT interval results, especially for the *SCN5A* gene, still illustrate the utility of this method. The availability of additional samples in the future will contribute to the success of this method in identifying genes of interest with greater statistical power. The gene annotation analyses revealed no clear BP-specific pathways or annotation, so these will also benefit from producing more specific and possibly larger gene sets. Additionally, we used hard genotype calls for analysis, necessitating some missing genotype data; the power of our methods could be improved by using imputed probabilities of genotypes.

Our attempts to expand findings beyond the known pathogenic coding variation with respect to the 20 genes involved in monogenic forms of hypertension or hypotension were inconclusive. We attribute this to the dearth of publicly available data for the kidney at this time, and expect that the availability of more extensive data will resolve some of the issues in further studies. Additionally, though it is beyond the scope of this study, as the effects of many of these monogenic disorders are likely through the adrenal gland, a full analysis of adrenal gland data will be necessary to assess them.

The MetaXcan software has supported most of the genes highlighted here and identified novel associations, although there were some limitations with the availability of the models for all genes. Additionally, our results indicated that deltaSVM weighting might be validly discriminatory between cell types; this is most evident with several QT interval genes, such as *NOS1AP* and *SCN5A*. It is also suggestive of cell-type specificity with the results for *CYP17A1* in the kidney cell types. It may be informative moving forward to characterize these BP genes at the individual cell-type level in the arteries as well.

The question of identification of core genes networks may be facilitated by our approach in this study, which includes using eQTL information from tissues or cell types of interest and genotypes to identify potentially relevant genes for a trait. As the expansion of publicly available resources continues, more information may be used for these purposes. Our analysis implicates specific variants that can be functionally tested for their effect on both gene expression and the phenotype.

## Materials and Methods

### Study participants and summary of genotypes, phenotypes, and association results used in this study

The full descriptions of the prior underlying studies, phenotypes, and association results for the GERA cohort are in Hoffmann et al. [38] and are briefly recapitulated here. The Genetic Epidemiology Research on Adult Health and Aging (GERA) cohort, part of the Kaiser Permanente Research Program on Genes, Environment, and Health (RPGEH), consists of individuals from five ethnic backgrounds; the majority is non-Hispanic white (EUR), with the remainder including Latino, East Asians, African Americans, and South Asians. A total of 99,785 individuals were analyzed, of which 80,792 were EUR individuals. The populations were each genotyped on custom population-specific Affymetrix Axiom SNP genotyping arrays [63, 64] and imputed to the 1000 Genomes Phase I Integrated Release Version 3 haplotype panel. Analyses of GERA alone, with the results of the International Consortium for Blood Pressure (ICBP, n=69,396) study [65], and with the ICBP and the UK Biobank (UKB, n=152,081) study [66], identified 316 novel BP loci. Combined with the set of replicated BP GWAS loci available at that time, there were a total of 390 BP loci we considered to be of interest. Of these, 367 had minor allele frequency (MAF) > 0.001 in the GERA EUR study, which was used as the reference population for the eQTL analyses described below.

For the purpose of several of the analyses described in this paper, we used these association results, as well as summary statistics available from 80,792 GERA EUR individuals from the Hoffmann et al. [38] study, and genotypes from a subset of 71,404 GERA EUR ‘unrelated’ individuals (third degree or beyond, pruned by the KING software for relationship inference) [67]. We converted genotypes prepared in the Hoffmann et al. [38] study after imputation from IMPUTE2 genotype probabilities format to PLINK ‘hard’ calls (the most likely genotype), setting genotypes with uncertainty greater than 0.25 to missing, and retaining variants with < 10% missing data, a Hardy Weinberg equilibrium test p < 1×10^-6^, and imputation quality score ≥ 0.3. In order to report univariate summary statistics within the 71,404 individuals, we used the --assoc option for analysis of a quantitative trait (Wald test) with PLINK v1.9 [68]. We analyzed covariate-adjusted longitudinal systolic (SBP) and diastolic (DBP) blood pressure in this study, as also described in Hoffmann et al. [38].

### ARIC genotypes, phenotypes, and association

The Atherosclerosis Risk in Communities (ARIC) study cohort is a longitudinal population-based study of 15,792 individuals, including 11,478 European-Americans (EUR) and 4,266 African-Americans (AA) from four study centers: Washington County, MD; Forsyth County, NC; Jackson, MS; and, Minneapolis, MN [24, 25]. The initial examination occurred from 1987-1989, with participants aged between 45 and 64 years. Subsequent examinations occurred in 1990-1992, 1993-1995, 1996-1998, and 2011-2013, with the most recent visits (6+) beginning in 2016. We analyzed 9,083 individuals of European ancestry with genotypes and QT interval at baseline. The genotyping of these samples on the Affymetrix genome-wide Human SNP Array 6.0, quality control, and imputation to the 1000 Genomes Phase I Integrated Release Version 3 haplotype panel are described elsewhere [69, 70]. We converted IMPUTE2 genotype probabilities to PLINK ‘hard’ calls, setting genotypes with uncertainty greater than 0.25 to missing, and retaining variants with < 10% missing data, a Hardy Weinberg equilibrium test p < 1×10^-6^, and imputation quality score ≥ 0.3, (as for the GERA study). The phenotypes were analyzed as previously described [71] with QT residuals generated by adjusting raw QT intervals for age, sex and resting heart rate. Summary statistics were generated for single variants using the --assoc option for analysis of a quantitative trait (Wald test) with PLINK v1.9 [68].

### GTEx genotypes and expression

We analyzed genotypes and expression data from the Genotype-Tissue Expression (GTEx; phs000424.v6.p1) Project [72] v6p for the SKAT analysis (see below) from the aorta, tibial artery, heart left ventricle, and heart atrial appendage tissues. Normalized expression was analyzed for these tissues, with the top three principal components, available PEER factors (15-35, depending on sample size), genotyping array platform, and sex used as covariates, all available from the GTEx portal. We used SNP-gene associations from the associated * .v6p.all_snpgene_pairs.txt.gz files from the authors’ eQTL analyses.

### Partitioned heritability analyses

We used the stratified LD score regression method and software [15] for estimating the heritability of the trait partitioned by genomic element using summary statistics for SBP and DBP from 80,792 GERA EUR individuals [38]. The ‘mungestats.py’ script was used to format the summary statistics as appropriate, and we analyzed them using the baseline model with 53 categories which included coding, UTR, and intronic regions, in addition to various open chromatin and histone modification annotations as described by the authors, as well as the 1000 Genomes Phase 3 reference files with the weights from their weights_hm3_no_hla.tgz file, which were provided and described by the authors on their website (https://github.com/bulik/ldsc/wiki/Partitioned-Heritability).

Generation of putative regulatory element maps and deltaSVM scores are described in the supplementary methods.

### Gene-based testing with SKAT

We used the sequence-kernel association test (SKAT) [22, 73] to test genes with median reads per kilobase of transcript, per million mapped reads (RPKM) ≥ 0.3 in GTEx samples for the aorta (n=197) and tibial (n=285) arteries with their respective variant sets. For each gene, we tested all variants within 50Kb of the gene start or end, inclusive of the entire gene body, per GENCODE v19 annotations (https://www.gencodegenes.org/releases/19.html). The weights used were taken as the absolute value of the deltaSVM score for each variant to reflect its predicted impact; for comparison, we also ran SKAT using default weights with beta density parameters (weights.beta=c(1,25), which up-weights rare variants as compared to common variants), as well as equal weights to all variants (weights.beta=c(1,1)). We tested association of each gene with adjusted SBP and DBP phenotype residuals (see above), as well as the GTEx normalized expression data with covariates (release v6p, https://www.gtexportal.org/), from the aorta and tibial arteries. We restricted our primary analyses in each of the kidney cell types to the 20 monogenic hypertension and hypotension genes. We additionally tested tissue- or cell-type-specific groupings in the ARIC dataset with the adjusted QT interval phenotype using the sets for the heart and heart tissues from GTEx, arteries and kidney cell types, as described above.

### Prediction of gene expression association with blood pressure

We used the MetaXcan [23] software with prebuilt HapMap training models for the GTEx (https://www.gtexportal.org/) tissues aorta and tibial arteries, provided by the authors at http://predictdb.hakyimlab.org/, with summary statistics from 80,792 GERA EUR individuals for SBP and DBP. We also used the software with the provided models for heart left ventricle and atrial appendage, for the QT interval analysis using summary statistics from 9,083 ARIC EUR individuals. MetaXcan is an extension of the PrediXcan [74] method, which predicts gene expression from genotypes and tests association of predicted expression with phenotypes using summary association results.

### Statistical significance

Statistical significance was determined using the Benjamini-Hochberg [75] (BH) method for multiple test correction to adjust for the number of genes within each analysis. We made no additional adjustments for the number of tissues, in part due to the correlation of specific subsets (the arteries, and individual kidney cell types), and as we examined genes across multiple analyses, for phenotype and for expression.

### Annotation of artery-significant genes

We used DAVID 6.8 [48, 49] to annotate the set of genes that met significance in the association analysis with either SBP or DBP. We retained terms that met a BH threshold of p < 0.05.

## Acknowledgements and Funding

The Atherosclerosis Risk in Communities Study is carried out as a collaborative study supported by National Heart, Lung, and Blood Institute contracts (HHSN268201100005C, HHSN268201100006C, HHSN268201100007C, HHSN268201100008C, HHSN268201100009C, HHSN268201100010C, HHSN268201100011C, and HHSN268201100012C), R01HL087641, R01HL59367 and R01HL086694; National Human Genome Research Institute contract U01HG004402; and National Institutes of Health contract HHSN268200625226C. The authors thank the staff and participants of the ARIC study for their important contributions. Infrastructure was partly supported by Grant Number UL1RR025005, a component of the National Institutes of Health and NIH Roadmap for Medical Research. The Genotype-Tissue Expression (GTEx) Project was supported by the Common Fund of the Office of the Director of the National Institutes of Health, and by NCI, NHGRI, NHLBI, NIDA, NIMH, and NINDS. The data used for the analyses described in this manuscript were obtained from: the GTEx Portal and dbGaP accession number phs000424.v6.p1. This research was funded by NIH grants HL128782 and HL0-86694 to A.C.

We are grateful to the Kaiser Permanente Northern California members who have generously agreed to participate in the Kaiser Permanente Research Program on Genes, Environment, and Health. Support for participant enrollment, survey completion, and biospecimen collection for the RPGEH was provided by the Robert Wood Johnson Foundation, the Wayne and Gladys Valley Foundation, the Ellison Medical Foundation, and Kaiser Permanente Community Benefit Programs. Genotyping of the GERA cohort was funded by a grant from the National Institute on Aging, National Institute of Mental Health, and the National Institute of Health Common Fund (RC2 AG036607 to CAS and NJR). GE receives support from Geneva University Hospitals and The Foundation of Medical Researchers, Geneva.

## Supporting Information

Fig S1. Comparisons of deltaSVM association P values between the heart and the other tissues in this study. hrt, heart.

Fig S2. Comparisons of -log10(P) differences of heart significant genes between deltaSVM weighting and equal weighting SKAT tests. hrt, heart; X, x-axis; Y, y-axis.

Fig S3. Aorta QQ plots, SBP.

Fig S4. Aorta QQ plots, DBP.

Fig S5. Tibial Artery QQ plots, SBP.

Fig S6. Tibial Artery QQ plots, DBP.

Fig S7. ENCSR000EOK QQ plots, SBP.

Fig S8. ENCSR000EOK QQ plots, DBP.

Fig S9. ENCSR000EOM QQ plots, SBP.

Fig S10. ENCSR000EOM QQ plots, DBP.

Fig S11. ENCSR000EPW QQ plots, SBP.

Fig S12. ENCSR000EPW QQ plots, DBP.

Fig S13. ENCSR785BDQ QQ plots, SBP.

Fig S14. ENCSR785BDQ QQ plots, DBP.

Table S1. Partitioned heritability results from baseline model for SBP.

Table S2. Partitioned heritability results from baseline model for DBP.

Table S3. deltaSVM performance results.

Table S4. SKAT and MetaXcan results for QT interval.

Table S5. Individual variants analyzed in kidney for C10orf32 and CYP17A1.

Table S6. DAVID enrichment analysis of artery-significant genes.

Text S1. Supplementary methods, results, and references.

