## Supplemental Text 1 for "Analysis of putative cis-regulatory elements regulating blood pressure variation"

**Supplementary Methods**

***Identifying genes at associated loci***

To identify potential target genes, we first determined the boundaries of associated loci using topologically associated domains (TADs) from published Hi-C data sets [1,2]. Specifically, we obtained full lists of contact domains identified by the Arrowhead algorithm on public Hi-C data sets for nine human cell lines (GEO accession no: GSE63525, GSE74072). Then, for each of the sentinel SNPs for QT interval and BP, the smallest domain containing the SNP was identified across all data sets. Genes that overlap any of the selected domains were identified as potential targets. We used the gene annotation from Gencode V19 (<https://www.gencodegenes.org/releases/19.html>). In total, we identified 94 genes for QT interval and 2,046 genes for BP variation.

***Construction of putative regulatory element maps***

Much of the statistical analyses described here required identification of putative enhancers that were active in tissues of interest. We constructed these genomic maps as follows. Putative regulatory element maps for the arteries were constructed using datasets available from ENCODE considered as representative of the aorta, tibial artery, and certain parts of the kidney (downloaded May 23, 2017). Open chromatin regions that were detected by either DNase-seq or ATAC-seq analyses were taken as putative regulatory elements; for the arteries, enhancer-associated H3K27ac or H3K4me1 histone modifications were used to subset those open chromatin regions that overlapped and were taken as putative enhancers. We completed this work as an extension of previous work in the generation of a cardiac CRE map [3].

As there were no whole aorta open chromatin datasets, we merged five artery open chromatin datasets for this tissue, available among the adult tissue datasets from ENCODE. These included DNase-seq datasets from two aortic cell types (smooth muscle (ENCSR000EIH), adventitia (ENCSR000EMC)) and two pulmonary artery cell types (endothelial cell (ENCSR000EOG) and fibroblast (ENCSR000EOH)), as well as one ATAC-seq dataset from whole tibial artery (ENCSR630REB). The tibial artery dataset was available in raw FASTQ format at the time of download, and was processed using the Kundaje lab protocol (<https://github.com/kundajelab/atac_dnase_pipelines>), with the exception of the BAM processing steps, including sorting, cleaning, duplicate marking and alignment filtering, retaining reads with quality score >30 and if properly paired, with samtools [4] v1.3 and Picard Tools [5] v2.9.2, were completed as described in (<https://github.com/tobiasrausch/ATACseq/blob/master/src/atac.sh>). We used five whole, ascending, or thoracic aorta H3K27ac datasets (ENCSR069UMW, ENCSR015GFK, ENCSR318HUC, ENCSR519CFV, and ENCSR322TJD).

For tibial artery, we used the tibial artery ATAC-seq dataset that was also used as a subset of the open chromatin regions for aorta above (ENCSR630REB), as well as one H3K4me1 dataset available for whole tibial artery (ENCSR233LCT). The peak calling for the H3K4me1 dataset was redone from the available filtered BAM file with MACS2 [6] with the --broad option to call broad peaks, using the parameters q=0.05 for the narrow peak cutoff, and q=0.2 for the broad peak cutoff, with extension size 400 bases.

While no whole kidney open chromatin data were available, there were four cell types from adult/non-fetal sources available in ENCODE: renal cortical epithelial cell (ENCSR000EOK), glomerular endothelial cell (ENCSR000EOM), epithelial cell of the proximal tubule (ENCSR000EPW), and glomerular visceral epithelial cell from a 3-year-old child (ENCSR785BDQ). No histone modification data were used, so open chromatin regions were considered as putative regulatory regions as a whole category.

For all experiments, where replicates were available, the one with the fewer peaks called was used as the reference replicate with peaks overlapping the replicate within each experiment retained. The artery open chromatin peaks were merged with the bedtools v2.26 merge utility [7]. To prepare the open chromatin regions for each tissue for training with gkm-SVM, we selected 600bp regions to represent each putative regulatory element. Peaks shorter than 600bp were extended 300bp from the midpoint, and longer peaks were the subset of the 600bp region with the maximum signal over all combined datasets for each tissue. The five H3K27ac datasets for aorta were merged using bedtools merge, and the peaks from this merged set and the tibial artery H3K4me1 dataset were extended by 100 bases in both directions, as we considered the boundaries of histone modification peaks to be inexact [8,9]. We retained those open chromatin regions overlapping a histone modification peak (after the 100bp extension) by at least 50% for training for the aorta and tibial arteries. Nearly all the datasets were available on genome build hg38, so these analyses were carried out on this build and then lifted over to hg19 for compatibility with downstream analyses.

***Generation of deltaSVM scores***

Each set of peaks (hg19 build) for the six tissues or cell types (aorta, tibial artery, and each of the four kidney cell types) was used as a positive training set to build gkm-SVM models [10], as previously described [10–12], with some modifications. The LS-GKM [13] software with the default parameter set was used for training. Regions from chromosome 9 were excluded from training and used for evaluating the adequacy of the trained models. The final models were generated by averaging gkm-SVMs from 10 independent negative training sets for each of the tissues or cell types. Subsequently, for each tissue or cell type, a deltaSVM score was calculated for each of 10,041,372 variants from the 1000 Genomes European-ancestry population, defined as the difference in gkm-SVM scores of the reference and alternate alleles for that variant, as previously described [14].

**Supplementary Results**

***Analysis of CREs in QT interval***

We analyzed the association of the QT interval adjusted for heart rate, age, and sex at baseline in 9,083 ARIC EUR individuals with putative CRE variants in genes at previous QT interval GWAS loci using SKAT and MetaXcan, with results depicted in Table S4. The tissues available from GTEx that we considered to be of relevance for QT interval for expression analyses are the heart left ventricle and atrial appendage tissues; for comparison, we also show the six tissues or cell types we analyzed in this study for blood pressure. We first found that several genes, including *SCN5A* and *NOS1AP*, showed significant association in heart tissues in our deltaSVM weighted SKAT CRE tests. Moreover, these associations are largely heart specific: comparisons of *P* values between the heart and the other tissues shows a systematic bias of p-values towards the heart (Fig S1). We also compared the -log_10_(P) differences of the heart significant genes between deltaSVM weighting and equal weighting SKAT tests and found more heart-specific effects (*i.e.* more genes with positive -log_10_(P) differences) (Fig S2). As specific examples, we further evaluated *SCN5A* and *NOS1AP*. A heart-specific effect is visible for *SCN5A* (deltaSVM p, p.SKAT.qt.dsvm =4.86 x 10^-4^; equal weighting p, p.SKAT.qt.eq=1.65 x 10^-4^) but not for the arteries or kidney cell types. Note that a much larger number of variants were analyzed for heart as there are more open chromatin regions within and immediately surrounding the gene in the heart compared to other tissues. The association with expression is not visible here, potentially due to a limited statistical power of eQTL analysis with small sample sizes. The *NOS1AP* gene shows a strong signal across all tissue/cell types, indicating the presence of individual variants driving the signal in open chromatin regions across all of the analyzed tissue/cell types. Interestingly, the well known and previously implicated QT interval signal, rs12143842, was only included in the heart variant set for analysis, and none of the other tissues or cell types, so these results are indicative of additional local contributions with higher functional weights. This said, we also observe an attenuation of signal by many orders of magnitude using deltaSVM weights compared to equal weights for many of the non-heart tissues; we see some evidence for this difference for *SCN5A* with the ENCSR000EOM results (p.SKAT.qt.dsvm=0.114; p.SKAT.qt.eq=0.0013). Additionally, it is also interesting to note that the SKAT analysis in GTEx heart left ventricle tissue shows the greatest evidence for association (p.SKAT.GTEx.dsvm=3.85 x 10^-4^; p.SKAT.GTEx.eq=1.51 x 10^-4^), with the next strongest signal from tibial artery (p.SKAT.GTEx.dsvm=8.67 x 10^-3^; p.SKAT.GTEx.eq=0.012). The MetaXcan results were uninformative in this case; there were no models available for *NOS1AP* in either heart tissue or for *SCN5A* in heart left ventricle (p.MetaXcan=0.011 in heart atrial appendage).
